# Long-read sequencing technology indicates genome-wide effects of non-B DNA on polymerization speed and error rate

**DOI:** 10.1101/237461

**Authors:** Wilfried M. Guiblet, Marzia A. Cremona, Monika Cechova, Robert S. Harris, Iva Kejnovska, Eduard Kejnovsky, Kristin Eckert, Francesca Chiaromonte, Kateryna D. Makova

**Author notes:** These authors contributed equally (Wilfried M. Guiblet, Marzia A. Cremona). Contact: Kateryna Makova 310 Wartik Lab Penn State University University Park, PA 16802, USA.

## Abstract

DNA conformation may deviate from the classical B-form in ~13% of the human genome. Non-B DNA regulates many cellular processes; however, its effects on DNA polymerization speed and accuracy have not been investigated genome-wide. Such an inquiry is critical for understanding neurological diseases and cancer genome instability. Here we present the first simultaneous examination of DNA polymerization kinetics and errors in the human genome sequenced with Single-Molecule-Real-Time technology. We show that polymerization speed differs between non-B and B-DNA: it decelerates at G-quadruplexes and fluctuates periodically at disease-causing tandem repeats. Analyzing polymerization kinetics profiles, we predict and validate experimentally non-B DNA formation for a novel motif. We demonstrate that several non-B motifs affect sequencing errors (e.g., G-quadruplexes increase error rates) and that sequencing errors are positively associated with polymerase slowdown. Finally, we show that highly divergent G4 motifs have pronounced polymerization slowdown and high sequencing error rates, suggesting similar mechanisms for sequencing errors and germline mutations.

## INTRODUCTION

The three-dimensional conformation of DNA at certain sequence motifs may deviate from the canonical double-stranded B-DNA (the right-handed helix with 10 nucleotides per turn)^1^ in helix orientation and strand number^2–4^. Approximately 13.2% of the human genome (394.2 Megabases) has the potential to form non-B DNA structures (Table S1), which are implicated in a myriad of cellular processes, and are associated with cancer and neurological diseases^3–7^. For instance, adjacent runs of guanines can form G-quadruplex (G4) structures^8^ (Fig. 1A) that participate in telomere maintenance^9^, replication initiation^10,11^, and transcriptional regulation^12^. Consequently, G4 structures have emerged as attractive anti-cancer therapeutic targets^13^. Additional non-B DNA structures associated with transcriptional regulation include left-handed Z-DNA duplexes formed within alternating purine-pyrimidine sequences^14^, A-phased repeats with helix bending formed within A-rich tracts^15^, and H-DNA triplexes formed within polypurine/polypyrimidine tracts and mirror repeats^16,17^ (Fig. 1A). Finally, Short Tandem Repeats (STRs), which also affect gene expression^18^, can adopt slipped-strand^19^ and other non-B DNA conformations^20^. Expansions of STRs are associated with numerous neurological and muscular degenerative diseases^21^. Notably, expansions of the hexanucleotide STR forming a G4 structure within the *C9orf72* gene is the most common genetic cause of amyotrophic lateral sclerosis (ALS)^22^. Moreover, STRs are enriched in cancer-related genes and participate in their functions^23^. Thus, growing evidence indicates that non-B DNA plays a pivotal role in several cellular pathways impacting health and disease.

**Figure 1.**
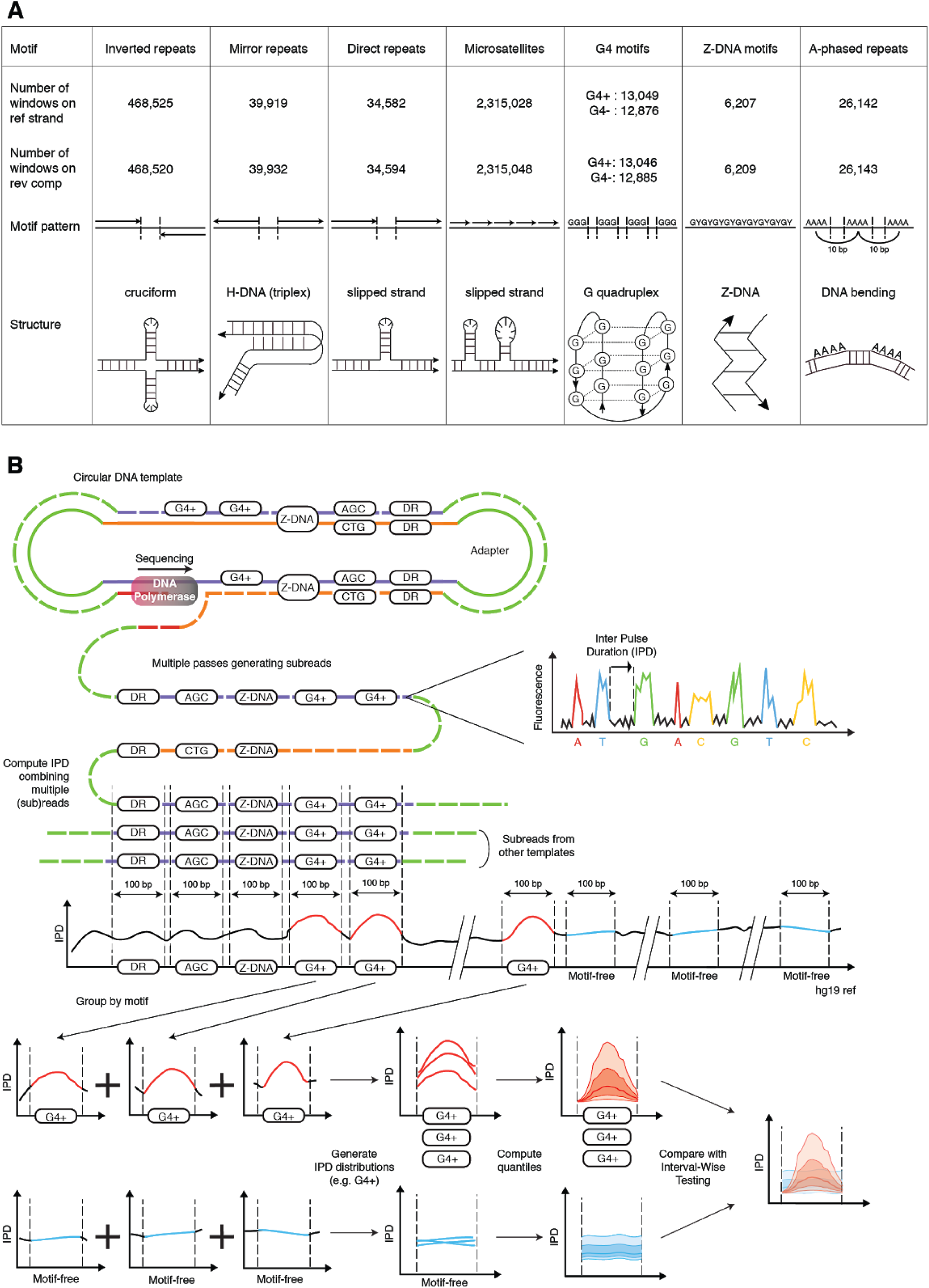
Non-B DNA motifs and Inter-Pulse Duration (IPD) analysis pipeline. **A** Non-B DNA motif types: patterns, putative structures, and counts of non-overlapping 100-bp windows containing one (and only one) motif with IPD measurements on the reference or reverse complement strand. **B** During SMRT sequencing, IPDs are recorded for each nucleotide in each subread (each pass on the circular DNA template). Subreads are aligned against the genome, and an IPD value is computed averaging ≥3 subread IPDs for each genome coordinate. We form 100-bp windows around annotated non-B motifs, extract their IPD values, and pool windows containing motifs of the same type to produce a distribution of IPD curves over the motif and its flanks (represented via 5^th^, 25^th^, 50^th^, 75^th^ and 95^th^ quantiles along the 100 window positions). We also form a set of non-overlapping 100-bp windows free from any known non-B motif, and pool them to produce a “motif-free” distribution of IPD curves. Each motif type is then compared to motif-free windows through Interval-Wise Testing (IWT). G4: G-quadruplexes; G4+/G4-: G4 annotated on the reference/reverse-complement strand; DR: direct repeats.

Whereas the transient ability of non-B DNA motifs to form non-canonical structures regulates many cellular processes^4^, these structures can also affect DNA synthesis and lead to genome instability, and thus can be viewed as both a blessing and a curse^24^. *In vitro* and *ex vivo* studies of individual loci showed that non-B DNA formation inhibits prokaryotic and eukaryotic DNA polymerases, causing their pausing and stalling of a replication fork^25–29^. These processes have been postulated to underlie non-B DNA-induced genome instability, i.e. increase in chromosomal rearrangements, including those observed in cancer^30,31^. Moreover, the increased occurrence of point mutations at non-B DNA was demonstrated at individual loci in plasmid constructs (reviewed in^4,32,33^), at disease-associated genes^34^, and among genetic variants from the 1,000 Genomes Project^35^. Because the effect of non-B DNA on mutagenesis is driven by both the inherent DNA sequence and polymerase fidelity^36^, we hypothesize that these structures can impact the efficiency and accuracy of DNA synthesis. Despite the critical importance of non-B DNA structures, ours is the first genome-wide study of their joint impact on polymerization speed and errors.

To evaluate whether DNA polymerization speed (i.e. polymerization kinetics) and polymerase errors are affected by non-B DNA, we utilized data from Single-Molecule Real Time (SMRT) sequencing. In addition to determining the primary nucleotide sequence, this technology, which uses an engineered bacteriophage phi29 polymerase^37^, records Inter-Pulse Durations (IPDs; Fig. 1B), i.e the times between two fluorescent pulses corresponding to the incorporation of two consecutive nucleotides^38^. We used IPDs as a measure of polymerization kinetics. Compared with other methods (e.g.,^39^), SMRT sequencing is presently the only high-throughput technology allowing a direct, simultaneous investigation of the genome-wide effects of several non-B DNA motif types on polymerization kinetics and errors. We also contrasted SMRT polymerization kinetics and sequencing error rates in highly mutable vs. invariant non-B DNA motifs, finding a potential link between polymerization in sequencing instruments and in living cells.

## RESULTS

### Non-B DNA motifs influence polymerization kinetics

We considered 92 different motif types potentially forming non-B DNA^4^ (Fig. 1A; Tables S1-S3), including predicted motifs from the non-B DNA DataBase^40^ and annotated STRs^41^. We constructed motif-containing genomic windows taking ±50 bp from the center of each motif (most were shorter than 100 bp; Fig. S1) and excluded overlapping windows (Tables S2-S3). For controls, we constructed 100-bp motif-free windows to represent genomic background, i.e. putative B-DNA. We populated each motif-containing and motif-free window with 100 single-nucleotide resolution IPDs (Fig. 1B) from a human genome previously sequenced with SMRT at 69×^42^. This was performed separately for the reference and reverse complement strands, because each strand is used separately as a template in SMRT sequencing (Fig. 1B). For each motif type, we aligned the centers of all motifs and aggregated IPD curves across windows, producing a distribution of IPD curves per strand (Fig. 1B).

To evaluate whether non-B motifs present polymerization kinetics patterns different from B-DNA, we used Interval-Wise Testing^43^, a novel Functional Data Analysis (FDA) approach, and identified genomic bases or intervals at which IPD curve distributions significantly differ between motif-containing and motif-free 100-bp windows (Fig. 2A-E; two-sided test, see Methods). We indeed found altered polymerization kinetics in and/or around several non-B DNA motifs. Below, we describe results for the reference strand (a total of 2,916,328 motif-containing and 2,524,489 motif-free windows; upper panels in Fig. 2A-D; Figs. 2E, S2-S10, S11A, S11C, S11E); results for the reverse complement serve as a biological replicate (lower panels in Fig. 2A-D; Figs. S11B, S11D, S11F).

**Figure 2.**
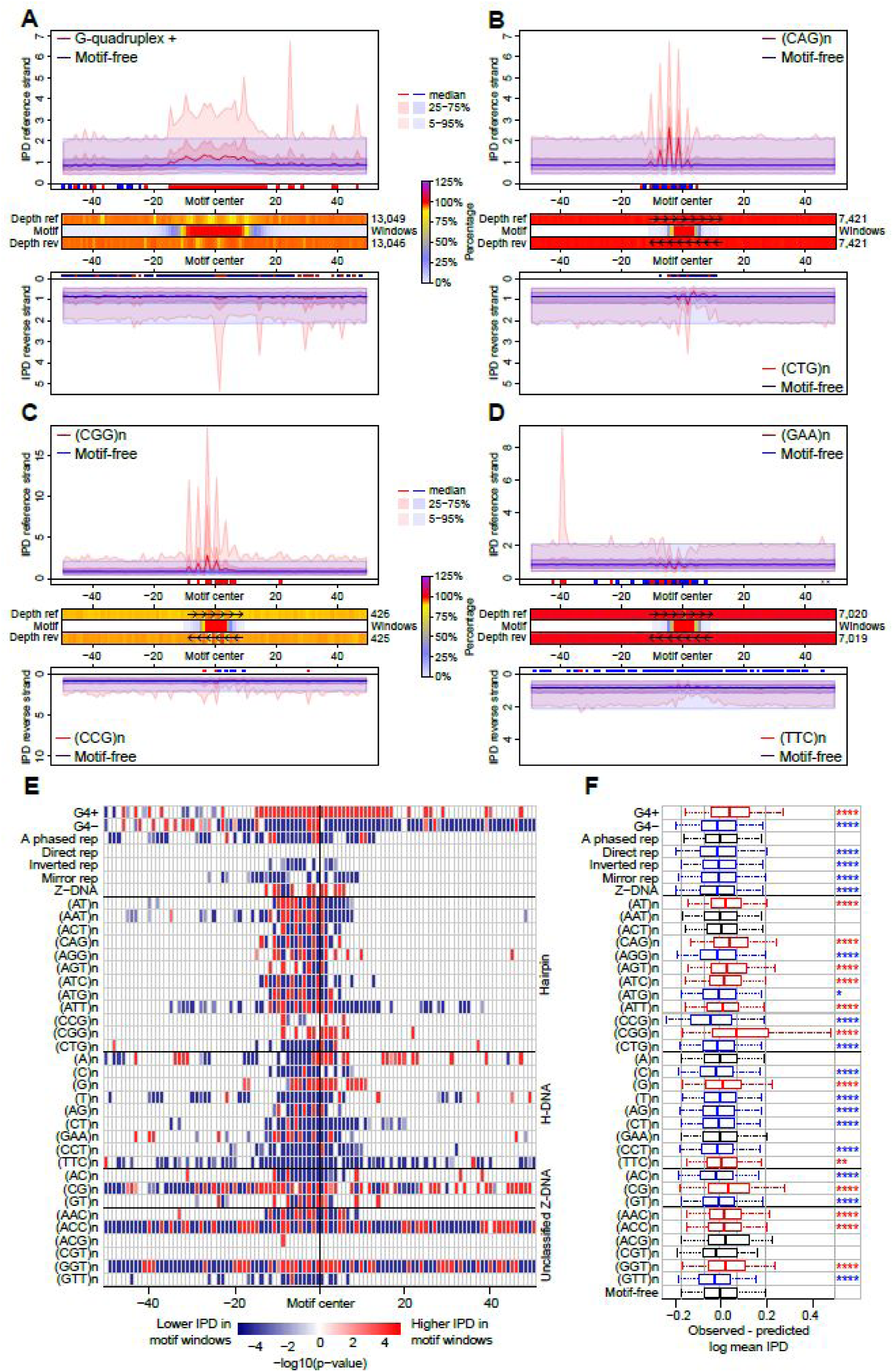
Polymerization kinetics at non-B DNA. **A-D** IPD curve distributions in motif-containing (red) vs. motif-free (blue) 100-bp windows, on reference (top) and reverse complement (bottom) strands. Thick lines: medians; dark-shaded areas: 25^th^-75^th^ quantiles; light-shaded areas: 5^th^-95^th^ quantiles. Red/blue marks (below top and above bottom plots): positions with IPDs in motif-containing windows higher/lower than in motif-free windows (IWT-corrected p-values≤0.05). Heatmaps (between top and bottom plots): sequencing depth of motif-containing relative to motif-free windows (in percentages, can be >100%) on reference (Depth ref) and reverse complement (Depth rev) strands, and percentage of windows with the motif (Motif) at each position. **A** G-quadruplexes; **B-D** STRs with disease-linked repeat number variation. **E** IWT results for IPD curve distributions in motif-containing vs. motif-free windows (reference strand). Each row shows significance levels (-log of corrected p-values) along 100 window positions for one motif type. White: non-significant (corrected p-value>0.05); red/blue: significant with IPDs in motif-containing windows higher/lower than in motif-free windows. STRs are grouped according to putative structure. **F** Comparison between observed mean IPDs in motif-containing windows and predictions from a dinucleotide compositional regression fitted on motif-free windows (reference strand). Bonferroni-corrected t-test p-values for differences: ≤0.0001 ‘****’, ≤0.001 ‘***’, ≤0.01 ‘**’, ≤0.05 ‘*’. Black: non-significant (corrected p-value>0.05); red/blue: significant with observed mean IPDs higher/lower than composition-based predictions. Boxplot whiskers: 5^th^ and 95^th^ quantiles of the differences.

Two lines of evidence are consistent with G4 motifs hindering polymerase progression. First, they decreased polymerization speed. Compared to motif-free windows, G4-containing windows had significantly higher IPDs near their centers, i.e. near the motifs (up to 1.7-fold IPD increase at the 95^th^ quantile; Fig. 2A). All G4 motif types exhibited this elevation, although the IPD curve shapes differed depending on the motif sequence (Fig. S3). Furthermore, the shape of the IPD distribution encompassing all G4 motif types remained the same (Fig. S4) when we limited our analysis to motifs forming the most stable G4 quadruplexes, as identified by *in vitro* ion concentration manipulations^39^. Second, sequencing depth was lower at G4 motifs than at motif-free windows (86% of motif-free depth; Fig. 2A), suggesting that the former hindered polymerization, resulting in fewer reads covering the motif. Polymerization slowdown and decreased sequencing depth were evident on the reference strand where G4s were annotated (upper panel in Fig. 2A), consistent with G-quadruplex structures forming only on the guanine-rich strand^5^. Elevated IPDs were observed in all sequencing passes through the same G4+ containing circular template (Figs. 1B and S5), suggesting that the structure is not resolved during sequencing. In contrast, the corresponding opposite strand (lower panel in Fig. 2A), as well as the reference strand where G4s were annotated on the reverse complement strand (“G4-” in Fig. 2E), — both cytosine-rich — showed a significant overall polymerization acceleration and displayed a smaller decrease in sequencing depth (92% of motif-free depth).

We observed that several other non-B DNA motifs, e.g., A-phased repeats, inverted repeats, mirror repeats, and Z-DNA, had significantly altered polymerization kinetics — both slower (higher IPD) and faster (lower IPD; Figs. 2E and S6). In contrast to the G4 motifs, the effects on polymerization kinetics were similar on the two sequenced strands (Figs. 2E and S11B) suggesting that, for these motifs, non-B DNA can be formed with similar probability on each strand during the sequencing reaction (Fig. 1B).

Additionally, we found that STRs altered polymerization kinetics in a length-and sequence-dependent manner (Figs. 2B-E, S7-S10); these variables impact the types and stability of non-B DNA structures that can form in addition to slipped structures (Table S4). For STRs with ≥2-nucleotide repeated units, the variation in polymerization kinetics was periodic, with the period (in bases) matching the length of the repeated unit, consistent with effects of strand slippage. This pattern was evident for trinucleotide STRs whose expansions at some loci are associated with neurological diseases (Fig. 2B-D), e.g., (CGG)_n_, (CAG)_n_, and (GAA)_n_ implicated in Fragile X syndrome, Huntington’s disease, and Friedreich’s ataxia, respectively^21^. Genome-wide, (CGG)_n_ repeats showed a strong periodic decrease in polymerization speed (elevated IPDs) on the annotated strand (up to 9-fold IPD increase at the 95^th^ quantile; Fig. 2C), consistent with their ability to form G4-like structures and hairpins^44^. The pattern for (CAG)_n_ repeats, also capable of forming hairpins^20^, was similar (Fig. 2B). Globally, STRs capable of forming hairpins (Table S4) presented the most striking polymerization deceleration and periodicity (Figs. 2B-C, 2E and S7). In contrast, STRs forming H-DNA (Table S4), including (GAA)_n_, accelerated polymerization (Figs. 2D-E and S8). For many STRs, significant deviations from background IPD levels were shifted 5’ to the annotated motif (Figs. 2E and S11), possibly due to polymerase stalling caused by difficulty in accommodating the alternative DNA structure within the polymerase active site.

The alterations in polymerization kinetics at non-B DNA motifs are not readily explained by either base modifications or by nucleotide composition. First, IPD patterns for most non-B DNA motifs were still clearly detectable in amplified DNA (Fig. S12), suggesting that they were not due to base modifications in the original template DNA^38^. Second, compositional regressions with either single nucleotide or dinucleotide composition explained only a relatively small portion of mean IPD variation among motif-free windows; 11.5% for single nucleotides and 20.8% for dinucleotides. Moreover, the mean IPDs in most motif-containing windows were significantly different from those predicted by such regressions (Fig. 2F and S13). Thus, nucleotide composition falls far short of explaining IPD variations at non-B DNA motifs (Figs. S13-S14). In particular, the mere presence of guanines in G4+ motifs cannot explain the overall substantial deceleration of polymerization observed at these sites.

### Polymerization kinetics and biophysical characteristics of G-quadruplexes correlate

To experimentally test whether non-B DNA structures can form at predicted motifs, we investigated the relationship between polymerization kinetics and biophysical characteristics of the ten G4 motifs most common in the human genome (Table S5). According to circular dichroism spectroscopy (CD) and native polyacrylamide gel electrophoresis (PAGE) analyses, all ten motifs quickly formed stable quadruplexes at low potassium concentrations, suggesting that they have a high propensity to form such structures^45^ albeit with different molecularity (intra-or intermolecular) and strand orientations (parallel or antiparallel, Table S5). Using regressions for intramolecular G4s, we found a significant positive relationship between mean IPD and delta epsilon (Fig. 3; p<2×10^-16^, R-squared=32.3%), a measure of structure organization quality obtained by CD^45^, and between mean IPD and melting temperature (solid line in cyan in Fig. S15B; p<2×10^-16^, R-squared=5.7%), a measure of thermostability and structure denaturation obtained by light absorption^45^ (Table S5; results for intermolecular G4s are shown in Fig. S15). Thus, polymerization slowdown and the biophysical characteristics of G4 formation are correlated, strongly suggesting that the motifs indeed form G4 structures during the SMRT sequencing reaction (intramolecular G4 structures are only a few nanometers in diameter^46^ and thus can fit within the 60×100 nm wells of Pacific Biosciences, or PacBio, instruments^47^).

**Figure 3.**
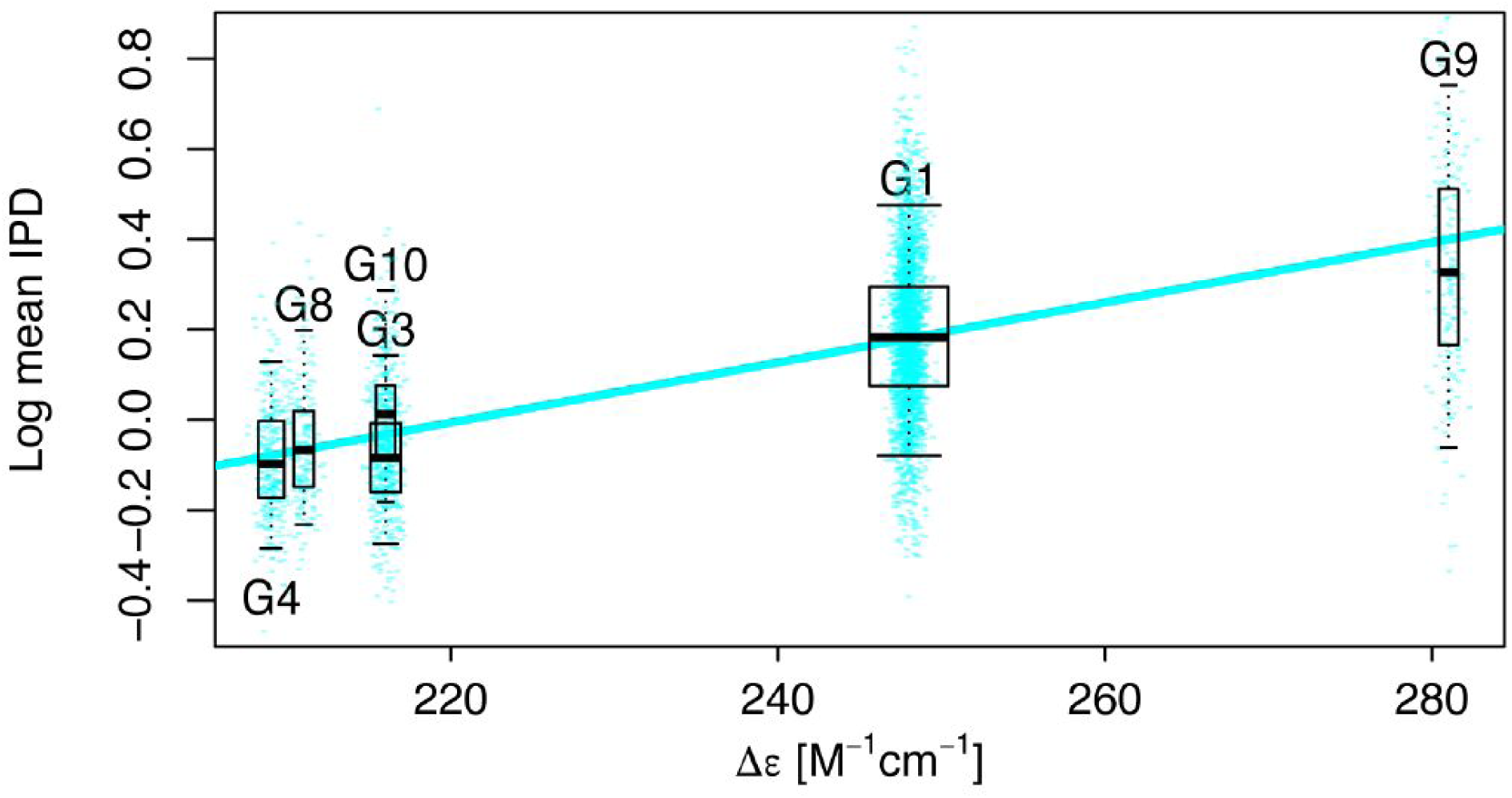
Relationship between G-quadruplex thermostability and polymerization kinetics. For the ten most common G-quadruplex motif types (G1 through G10, in order), we measured circular dichroism (delta epsilon) and light absorption (melting temperature, T_m_), and computed average IPD values for each of thousands of motif occurrences in the genome (Table S5). For intramolecular G4s, average IPDs were regressed on delta epsilon (R-squared = 32.3%). Boxplot whiskers: 5^th^ and 95^th^ quantiles. Boxplot width: proportional to the square root of the sample size for each motif. Points: individual occurrences used in the regressions, with horizontal jittering for visualization (results for delta epsilon in intermolecular G4s and for T_m_ are shown in Fig. S15).

Our experiments also showed that statistical FDA techniques applied to polymerization kinetics data can enable non-B DNA structure discovery. While not possessing a canonical G4 motif, the (GGT)_n_ STR has an IPD profile similar to that of G4+ (Figs. 2E and S10E) and its reverse complement (ACC)_n_ has an IPD profile similar to that of G4- (Figs. 2E and S10B), suggesting that (GGT)_n_ may fold into a G4-like structure. Remarkably, biophysical analyses (CD, native PAGE, and thermal denaturation) showed that (GGT)_n_ motifs indeed adopt quadruplex conformation (Fig. S16; Table S6).

### Non-B DNA motifs affect sequencing error rates

To examine whether phi29 polymerase accuracy is affected during synthesis of non-B DNA motifs of different types in the genome, we contrasted SMRT sequencing error rates between such motifs and motif-free regions. Error rates were computed using the same human genome sequenced at 69×^42^ that was used for the IPD analysis above. Because of the potential for inaccurate typing of STRs^41^ and for motif misalignments in repetitive loci, we restricted our attention to six non-STR motif types present on the reference strand of the non-repetitive portion of the genome (Tables 1 and S7). We focused on motifs themselves (as opposed to 100-bp motif-containing windows), and for controls we identified motif-free regions matched to motifs in number and length. We excluded motifs and moti-free regions with fixed differences between sequenced and reference genomes (i.e. with germline variants present in the sequenced genome compared to the reference), and computed sequencing error rates as the proportion of variants (relative to hg19) within the total number of nucleotides sequenced for the motif or motif-free region — including errors supported even by a single read (see Methods). Because we were interested in a detailed analysis of errors made by the polymerase, we used raw SMRT reads and not the circular consensus sequences. Below we present results for errors on the newly synthesized strand that uses the template strand annotated with non-B DNA motifs.

We observed a strong effect of G4 motifs on SMRT error rates. Mismatches were markedly elevated (1.79-fold) on the newly synthesized strand when G4s were present on the template strand (Table 1). Deletions were increased in both G4+ and G4- (Table 1; 1.49-and 1.11-fold, respectively). Insertions, the most common error type for SMRT sequencing, were depressed when the templates encoded G4+ and particularly G4-motifs (Table 1). In contrast to G4 motifs, Z-DNA displayed depressed mismatches and deletions, but increased insertions (Table 1). In summary, the rates of all three types of SMRT sequencing errors differed between non-B motifs and motif-free regions, with strong elevations of mismatches and deletions at G4-motifs.

**Table 1.** Error rates at non-B DNA motifs during SMRT sequencing. We contrasted rates of SMRT sequencing errors between the indicated motifs and motif-free regions. Numbers are fold differences; significance is indicated by asterisks (p≤0.0001 ‘****’, ≤0.001 ‘***’, ≤0.01 ‘**’, ≤0.05 ‘*’, ≤0.10 ‘.’). Significant values are shown in bold. Sample sizes are in Table S7.

| SMRT errors | A-Phased Repeats | Direct Repeats | Inverted Repeats | Mirror Repeats | Z-DNA | G4+ | G4- |
| --- | --- | --- | --- | --- | --- | --- | --- |
| Mismatches | -1.018 | **1.089**<br>**** | 1.004 | **1.002**<br>** | **-1.192**<br>**** | **1.790**<br>**** | 1.058 |
| Insertions | **-1.032**<br>**** | -1.016<br>. | **-1.020**<br>**** | **-1.023**<br>**** | **1.156**<br>**** | **-1.037**<br>**** | **-1.230**<br>**** |
| Deletions | **-1.042**<br>**** | **1.018**<br>**** | **-1.000**<br>**** | 1.001 | **-1.150**<br>**** | **1.494**<br>**** | **1.106**<br>**** |

We next tested whether SMRT mismatch error rates could be explained by sequence composition. Only 4.1% of the variability in SMRT error rates in motif-free windows could be explained by single nucleotide composition (Fig. S17A). Among the four nucleotides, the content of guanines was the most correlated to SMRT errors, and its increase led to elevated SMRT error rates (Fig. S17A). Dinucleotide compositional regression also explained a rather small proportion of variability in SMRT error rates in motif-free windows (R-squared=5.6%). Furthermore, SMRT error rates in most types of motifs (in all but A-phased repeats) were significantly different than those predicted by such compositional regressions (Fig. S17B). Thus, nucleotide composition does not suffice to explain SMRT error rate variation at both motif-free windows and non-B DNA motifs. In particular, the high concentration of guanines in G4+ motifs cannot explain the increase in SMRT error rates observed at such sites.

### Increased SMRT error rates are associated with polymerase slowdown, particularly at non-B DNA

We next studied whether SMRT error rates are associated with polymerization speed. We focused on G4+ and G4-motifs, which had the strongest effect on SMRT error rates among the non-B motif types examined (Table 1), and used motif-free windows for comparison. We fitted a regression expressing SMRT mismatch error rates as a function of mean IPD values corrected for nucleotide composition (residual mean IPDs, i.e. the difference between observed mean IPDs and the ones predicted using single nucleotide composition, see Methods). The model also accounted for the three groups of regions — G4+ motifs, G4-motifs, and motif-free windows — and had an overall R-squared of 35.4% (Fig. 4). We found a significantly positive linear relationship between SMRT mismatch rates and residual mean IPDs in motif-free windows (slope=0.11, p=2.9×10^-10^). Interestingly, the slope of the regression line was significantly steeper for G4+ than for motif-free windows (slope=0.26, p=1.9×10^-12^ for difference with motif-free), while G4- had a slope similar to motif-free windows (slope=0.08, p=0.17 for difference with motif-free). Thus, SMRT mismatch errors are positively associated with polymerase slowdown, and this association is particularly strong in G4+.

**Figure 4.**
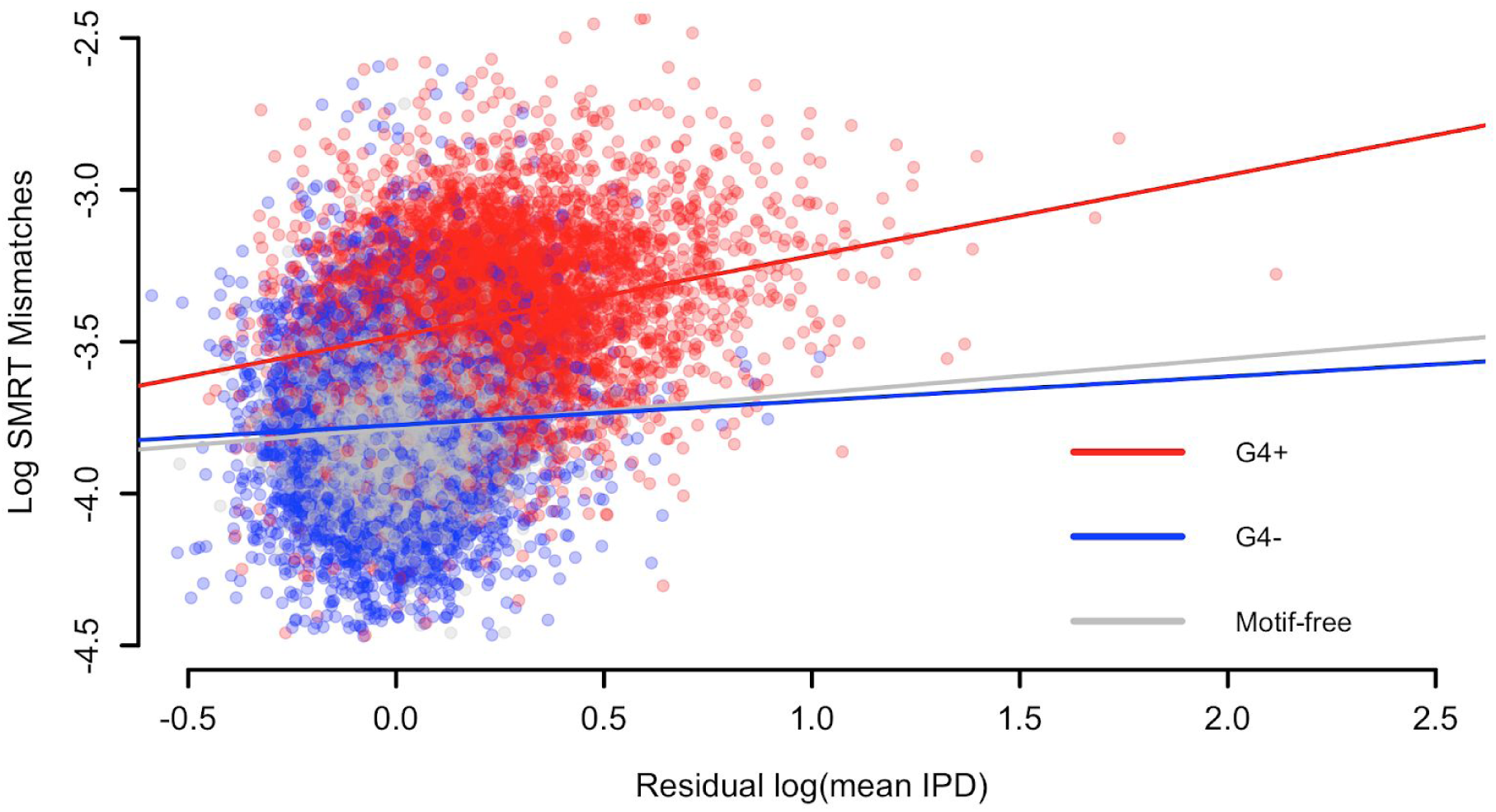
Errors are linked to kinetic variation. Sequencing through templates containing G-quadruplex motifs (G4+) increases the positive relationship between mismatches in SMRT sequencing and IPD values (corrected for sequence composition). This does not occur when sequencing through the reverse complement of G-quadruplex motifs (G4-). The regression model accounting for three groups of regions (G4+, G4-, and motif-free windows) has R-squared equal to 35.4%. Slopes are 0.26, 0.08, and 0.11 for G4+ motifs (N=5,937), G4-motifs (N=5,695), and motif-free windows (N=11,632, which is the sum of samples sizes for G4+ and G4-), respectively.

### Polymerase speed and mutation occurrence

Mutation rates are known to be non-uniform across the genome; however, the mechanisms leading to such regional variation are not entirely understood^48^. Our results on sequencing errors for the SMRT technology, as well as previous *in vitro* polymerase studies^27,28,49,50^ demonstrating the effects of non-B DNA on DNA synthesis by phage, prokaryotic and eukaryotic polymerases, raise an intriguing question: are mutation rates *in vivo* also affected by these motifs via polymerase slowdown? Environmental influences apart, mutations are the net result of polymerase errors and the lack of repair in the cell. Here we are making a simplifying assumption that mutations result primarily from polymerase errors^51^. Examining the speed of polymerases at the nucleotide level in eukaryotic cells is a challenging endeavor, but can the polymerization kinetics and error profile of the phi29 enzyme provide us with a hint of how non-B DNA motifs might be affecting mutations? To address this question, we contrasted SMRT error rates and mean IPDs between G4+ motifs with high and low human germline mutation rates approximated by the level of human-orangutan divergence or by the level of intraspecific diversity inferred from the 1,000 Genomes Project^52^. In more detail, we compared SMRT mismatch error rates and mean IPDs (corrected for single nucleotide composition, see above) between G4+ motifs in the top 5% vs. in the bottom 5% of human-orangutan divergence, as well as between G4+ motifs in the top 5% vs. in the bottom 5% of diversity from the 1,000 Genomes Project (Table S9). Human and orangutan genomes^53,54^ were generated with the Sanger technology, and only common high-frequency variants (with minor allele frequency equal to or above 1%) were considered for the diversity analysis. Therefore, divergence and diversity data are expected to be highly accurate.

Highly divergent (or diverse) G4+ motifs had higher IPD values (i.e. experienced polymerase slowdown), as compared with G4+ motifs having low divergence (or diversity) (p=4×10^-04^ and p=0.046 for divergence and diversity, respectively; t-test for difference in means; Table S9). Moreover, highly divergent (or diverse) G4+ motifs had higher error rates than did G4+ motifs with low divergence (or diversity; p=0.040 and p=0.014 for divergence and diversity, respectively; t-test for difference in means; Table S9). Therefore, indeed, we found divergence (diversity) to be negatively related to polymerization speed and positively related to SMRT sequencing errors, suggesting that G-quadruplexes affect not only sequencing errors, but also germline mutations *in vivo*.

## DISCUSSION

SMRT sequencing polymerization kinetics data are produced during every SMRT sequencing experiment, but are rarely analyzed by researchers, except for studies of DNA modifications (e.g., ^55^). Our genome-wide study exemplifies the usefulness of such data in four additional scientific domains: (1) studies of polymerization kinetics; (2) discovery of novel non-B DNA structures; (3) analysis of sequencing errors; and (4) correlating polymerase kinetics with error rate. With the increasing popularity of SMRT sequencing and growing publicly available data, we expect an acceleration of progress in these areas.

Analyzing hundreds of thousands of non-B DNA motif occurrences genome-wide, we observed polymerization slowdown or acceleration during SMRT sequencing for the majority of motifs types considered. Particularly striking patterns were noted for G4s and STRs, for which we found strong slowdown and periodic alterations in polymerization speed, respectively. Our study corroborates a previous analysis of 27 occurrences of (CGG)_n_ — a motif capable of forming a G4 structure and a hairpin — which indicated polymerization speed alterations in the *E. coli* genome^56^. It was suggested that non-B DNA act as a polymerization speed modifier along the genome not only in a sequencer, but also under natural conditions, i.e. in the cell^56^. Backing of this hypothesis also comes from *ex vivo* analyses of a handful of loci capable of non-B DNA formation^25,26^. Future experiments should examine this intriguing possibility on a genome-wide basis. Also, our results lend support to the hypothesis that periodic polymerization kinetics patterns at disease-associated STRs contribute to their instability^57^.

We demonstrate that analyzing polymerization kinetics data with FDA statistical techniques can enable discovery of novel motifs forming non-B DNA structures. We identified (GGT)_n_ motifs as potentially forming a G4-like structure based on their polymerization speed patterns, and used biophysical profiling to validate the formation of such structure. The genomic distribution and potential function of such newly identified non-B motifs should be studied further. Moreover, our study shows that statistical analyses of IPD patterns can characterize non-B DNA in a way that is orthogonal to conventional biophysical profiling.

Elevated SMRT sequencing errors in many non-B DNA motifs should be taken into account when evaluating sequencing results. Many such errors are likely corrected during circular sequencing, but biases might remain in the sequence consensus and when using the long sequencing read mode (as opposed to the circular consensus sequencing mode). Of particular interest is our result concerning the non-randomness of insertions, which are the most common type of SMRT errors. We observe that insertions increase in Z-DNA, but in fact decrease in other non-B DNA motif types. In light of this, re-calibrating base quality scores in SMRT sequencing reads should be considered in future work.

To our knowledge, our analyses are the first to document an association between polymerization speed and accuracy on a genome-wide scale. We find that the phi29 polymerase makes more errors when the IPD is high, i.e. when polymerization is slowed down. This effect is significant for B-DNA, and further magnified in G-quadruplexes, even after correcting for nucleotide composition. Thus, polymerase accuracy may be affected by DNA sequence and structural features that hinder processive synthesis. Consistent with our findings here, previous *in vitro* studies showed a positive correlation between DNA polymerase error rates and pausing during synthesis (e.g., slow synthesis). Notably, misincorporation errors by the replicative DNA polymerase alpha are associated with polymerase pausing^58^, and polymerase alpha errors within STRs are positively correlated with polymerase pausing and non-B DNA formation^59^.

Our study indicates a significant effect of non-B DNA on the fidelity of DNA synthesis. With a genome-wide analysis of data generated by a sequencing instrument, we substantially expanded prior knowledge of this phenomenon gained by examination of plasmid constructs, disease-associated genes, and human genetic diversity ^4,32,33,35^. Our most prominent observation, the high incidence of mismatches and deletion errors during sequencing of G4 motifs, is in line with these motifs harboring excessive disease-causing point mutations^34^ and nucleotide variants (point mutations and indels combined^35^) based on 1,000 Genomes Project data. Using the level of divergence (or diversity) as a proxy for germline mutation rates, we observed significantly larger SMRT polymerase slowdown and error rates within G4s with very high divergence as compared to G4s with very low divergence (diversity). Selection is expected to decrease divergence and diversity levels, however we have no reason to predict that G4s undergoing stronger selection should have faster rates of DNA synthesis and lower sequencing error rates. Thus a link between polymerase accuracy and germline mutation rates is more plausible. Our results, taken together with previously published studies, argue for the pervasive role of G4s in affecting polymerase fidelity and germline mutation rates in the genome. Future experiments are needed to determine whether non-B DNA motifs affect polymerase kinetics and errors within living cells.

Compared with errors in sequencing instruments, mutations in the cell are the net result of DNA synthesis errors by more than 15 different polymerases and enzymes from numerous DNA repair pathways^60^. Furthermore, mutation occurrence is influenced by additional factors (e.g. chromatin, etc.^61^). Notwithstanding all these caveats, our results, together with published results of Du and colleagues^35^, lend support to the notion that non-B DNA is one among the “local DNA environment” factors affecting mutation rates and patterns^62^. Future studies should specifically investigate what share of local variation in mutation rates along the genome can be explained by the presence of non-B DNA. Our findings, together with observations on the transient nature of non-B DNA conformations^4,63,64^, portray non-B DNA as effective modulator of genome structure. This is particularly significant in view of recent evidence broadening the spectrum of mechanisms through which non-B DNA may modulate the cell, encompassing, e.g., epigenetic instability^24^ and non-coding RNA regulation^65^.

## METHODS

### Non-B DB and STR annotations

Annotations of A-phased, direct, inverted and mirror repeats, G-quadruplexes and Z-DNA motifs were downloaded from the non-B DataBase (DB) at https://nonb-abcc.ncifcrf.gov. Additionally, we annotated STRs on the human reference (hg19) using STR-FM^41^. We only considered mono-, di-, tri-, and tetranucleotide STRs with ≥8, ≥4, ≥3, and ≥3 repeats, respectively^41^. We then collapsed STR motifs that could be matched by changing their reading frame (Table S9). For instance, (AGC)_n_, (CAG)_n_, and (GCA)_n_ were collapsed into the (AGC)_n_ group. We restricted our attention to non-B motifs and STRs annotated on autosomes.

### Constructing genomic windows

Polymerization kinetics was studied in 100-bp windows (Fig. 1B). Motif-containing windows were centered at the middle coordinates of the annotated motifs in our list (Fig. S1). The centers of STRs with different repeat numbers were shifted to ensure their alignment (Table S8). Overlapping motif-containing windows (with motifs of the same or different type) were filtered out, leaving a total of 2,926,560 windows. All windows not containing motifs and not overlapping motif-containing windows were labeled as motif-free (a total of 3,649,152 windows).

### IPDs

We used publicly available PacBio resequencing data (69x) from an individual male (HG002; NA24385) belonging to the Genome in a Bottle Ashkenazim trio^42^. We analyzed 228 SMRT cells sequenced with P6-C4 chemistry in a mode maximizing the subread length and not the number of passes (analysis of the sequences obtained with P5-C3 led to similar results; data not shown)^66^. On average, each molecule was sequenced in 2.12 passes, with the majority of the molecules sequenced only in a single pass resulting in a single subread (74.76% of the molecules). Sequencing reads were aligned to hg19 with pbalign (smrtanalysis-2.3.0), resulting in an ~52x average read depth, and IPDs were computed at nucleotide resolution with *ipdSummary.py* (https://github.com/PacificBiosciences) — this produces one IPD value per site averaging among at least three subreads, normalizing for inter-molecule variability and trimming for outliers. The resulting IPDs, which are strand-specific (any observed slowdown or acceleration of the polymerization concerns the strand used as template), were then used to populate motif-containing and motif-free 100-bp windows according to their coordinates (Fig. 1B); each window thus contains an IPD curve comprising 100 values or less (if some nucleotides lack IPDs). All windows with no IPD values were filtered out, and only motifs with ≥15 windows with IPDs on both strands were retained for subsequent analyses. This left us with a total of 2,916,328 motif-containing and 2,524,489 motif-free windows on the reference strand, and 2,916,377 motif-containing and 2,524,612 motif-free windows on the reverse complement strand. Next, for each motif type (Tables S2-S3), and separately for each strand, we aligned the 100-bp windows. This resulted in strand-specific IPD curve distributions for each motif type. An IPD curve distribution was visualized plotting quantiles (5^th^, 25^th^, 50^th^, 75^th^ and 95^th^) of the IPD values at each of the 100 nucleotides along the aligned windows (see Figs. 1B, 2A-D, S3-S4, S6-S10). IPD distributions were visually unaffected by variants between the sequenced and the reference genomes.

### Interval-Wise Testing for differences in IPDs

To detect statistically significant differences between IPD curve distributions in motif-containing and motif-free windows, separately for each motif type and strand, we employed the Interval-Wise Testing (IWT) procedure for “omics” data implemented in the R Bioconductor package and Galaxy tool *IWTomics*^43,67,68^. IWT treats the IPD values in a 100-bp window as a curve (see Fig. 1B) and assesses differences between two groups of curves (containing a given motif, and motif-free) performing a non-parametric (permutation) test at all possible scales, from the individual nucleotides to the whole 100-bp. When IWT detects a significant difference at a particular scale, it also identifies the locations (window coordinates) that lead to the rejection of the null hypothesis (see Supplementary Text for details). Because IWT is computationally expensive, we ran it on a maximum of 10,000 curves for each motif type and strand (sample sizes are listed in Tables S2-S3). For motif types with *n* ≥10,000 windows, we randomly subsampled 10,000 windows and tested against a random set of 10,000 motif-free windows; this was repeated 10 times to ensure results robustness. For motif types with *n* ≤10,000 windows, we tested both against a random set of 10,000 motif-free windows and against a random set of *n* motif-free windows; in both cases we repeated the comparison against 10 random sets, again to ensure results robustness. IWT was performed using three test statistics: the mean difference, the median difference, and the multi-quantile difference (i.e. the sum of the 5^th^, 25^th^, 50^th^, 75^th^ and 95^th^ quantile differences). Results for the latter, which most effectively captures differences in curve distributions, are presented in Figs. 2E and S11A-B, while those for mean and median are presented in Figs. S11C-D and S11E-F, respectively. P-values were computed using 10,000 random permutations (independent samples, two-tailed test). The procedure produces an adjusted p-value curve (comprising 100 p-values, one for each nucleotide, adjusted up to the selected scale) for each comparison (Fig. S2). We summarized results for all motif types in adjusted p-value heat maps (Figs. 2E and S11, multi-quantile difference; Fig. S11, mean and median). Red/blue indicate positive/negative observed differences, and are shown only for significant locations (adjusted p-value ≤0.05 in each of the 10 repetitions).

### Effect of sequence composition on IPDs

To investigate whether differences in IPD values depend on incorporation of different nucleotides, we computed mean IPD, a single nucleotide composition vector *P_Si_* =(*p_A_*, *p_T_*, *p_C_*, *p_G_*) (*p_A_* + *p_T_* + *p_C_* + *p_G_* = 100%), and a dinucleotide composition vector *P_Di_* =(*p_AA_*, *p_AC_*, *p_AG_*, …, *p_TT_*) (*p_AA_* + *p_AC_* + *p_AG_* + … + *p_TT_* = 100%) in each 100-bp window. We considered only motif-free windows and combined data from both strands (results from the two strands considered separately were similar; data not shown). First, we measured the marginal effect of each nucleotide *j=A,C,G,T* as the correlation between *log*(*mean IPD*) and *p_j_*. Next, we employed compositional regression models^69,70^ to quantitate the overall effect of single nucleotide and dinucleotide composition on IPDs. The single nucleotide sequence composition vector *P_Si_* was mapped to a three-dimensional euclidian vector *X_Si_* =(*x*_1_, *x*_2_, *x*_3_) using the isometric log-ratio transform, and a multiple regression model was fitted for *log*(*mean IPD*) on *x*_1_, *x*_2_, *x*_3_. Model assumptions and validity were checked with standard multiple regression diagnostic plots and tests, and the R-squared was used to evaluate composition effect strength. Similarly, the dinucleotide composition vector was mapped to a 15-dimensional *P_Di_* euclidian vector *X_Di_* = (*x*_1_, *x*_2_, …, *x*_15_), and a multiple regression model was fitted for *log*(*mean IPD*) on *x*_1_, *x*_2_, …, *x*_15_. The dinucleotide compositional regression model fitted on motif-free windows, which had higher R-squared (see Results), was then used to predict the mean IPD values of motif-containing windows based on their composition, separately on each strand. For each motif type, we computed the differences between these predictions and observed mean IPDs (on logarithmic scale), created their boxplots and performed two-sided t-tests for the mean difference being equal to zero — using a Bonferroni correction to adjust for multiple motif testing (Figs. 2F and S13).

### Experimental characterization of G-quadruplexes

The ten most common G-quadruplex motifs (Table S5) from non-B DB annotations, as well as the (GGT)_n_ motifs, were studied by circular dichroism (CD), native polyacrylamide gel electrophoresis (PAGE) and UV absorption melting profiles, as described previously^71^. Single-stranded oligos were used in structure characterization of G-quadruplexes, for three reasons. First, G-quadruplexes often play a regulatory role in molecular processes where DNA is single-stranded, such as replication, transcription, and repair^72^. Second, single-strandedness allows better characterization of quadruplex formation and thus is most often used in experimental studies^73^. And third, the analysis of single-stranded structures is most relevant for SMRT sequencing, where single-stranded DNA molecules are used as templates.

Initially, we considered only intramolecular G-quadruplexes, computed the mean IPD in each occurrence of the motifs, and fitted a simple regression for *log*(*mean IPD*) on delta epsilon (for each motif delta epsilon was measured once, while mean IPD was computed for hundreds or thousands of occurrences; Table S5 and Figs. 3 and S15A). Next, we considered both intra-and intermolecular G-quadruplexes, and fitted a multiple regression for *log*(*mean IPD*) on delta epsilon, the molecularity of the G-quadruplexes (either intra or intermolecular; a binary predictor), and their interaction. We fitted similar single and multiple regressions (considering only intramolecular G-quadruplexes, and both intra-and intermolecular G-quadruplexes) replacing delta epsilon with melting temperature (T_m_; Fig. S15). In both cases we identified final models with backward selection.

### SMRT sequencing errors

Data are again those from PacBio sequencing of HG002; NA24385^42^. Errors were analyzed restricting attention to motif occurrences (*not* motif-containing 100-bp windows). Due to potential misalignments at motifs in the repetitive parts of the genome, motifs and motif-free windows overlapping with RepeatMasker annotations (rmsk track obtained at https://genome.ucsc.edu) were excluded from this analysis. To focus on errors and not on fixed differences, all motifs and motif-free windows overlapping variants between HG002 and hg19 were also excluded (we used high confidence calls from a benchmarking dataset generated in^42^). For each motif type, control sets were constructed picking a filtered motif-free 100-bp window at random from within 0.5 Mb upstream or downstream of each motif occurrence, and trimming it to produce a motif-free region of the same length of the motif occurrence itself. This matches motif occurrences and motif-free regions in number and length (which guarantees the same measurement resolution for errors), as well as in broad genomic location. We note that results are virtually unchanged if we do *not* match broad genomic location and select controls completely at random from the genome.

Error rates (the number of mismatches, insertions or deletions relative to hg19, divided by the total number of nucleotides from all subreads in a given region and expressed as a percentage) were calculated for the newly synthesized strand that used six non-STR motif types and corresponding motif-free regions as a template. Since our purpose was detecting polymerase errors, we calculated the error rates based on individual subreads by accessing the alignment files directly and considering also low-frequency variants, including those supported by a single subread.

### Comparison of errors between motifs and motif-free regions

To compare error rates between motif occurrences and matching motif-free regions, we employed a two-part test^74,75^ that contrasts both the heights of spikes at error rate 0 (corresponding to regions without errors) and the distributions of positive error rates. The compound null hypothesis is that both the spike at 0 (proportion of 0 rates) and the distribution on positive values (continuous component on non-0 rates) are the same in the two groups, versus the two-sided alternative that either or both differ between the groups. We considered the two-part statistic *V*^2^ = *B*^2^ + *T*^2^, where *B*^2^ is the continuity-corrected binomial test statistic (contrasting the proportions of 0 rates) and *T* ^2^ is the square of the t-test statistic (contrasting the non-0 rates). P-values were generated approximating the distribution of the test statistic *V*^2^ under the null hypothesis with a χ^2^(2), in order to overcome the computational burden of estimating its distribution using permutations. For several cases, we also computed p-values based on 10,000 random permutations and obtained almost indistinguishable results. For robustness, each test was repeated 10 times, using separate sets of randomly generated matching motif-free regions, and significance was assessed based on the maximum p-value (maximum p-values ≤0.10 are coded by standard stars-and-dots representation in Table 1).

In addition to running the tests, we computed rate fold differences (the numbers in Table 1) as follows. For each motif type, we considered the whole portion of the genome covered by its occurrences. For comparison, we considered the portion of the genome covered by *all* non-repetitive, without fixed differences 100-bp motif-free windows (note: *not* matching motif-free regions). SMRT error rates were estimated dividing the total number of errors by the total number of bases sequenced in the portion of the genome under consideration. Rate fold differences were then computed, for each motif type and each error type, as motif rate over motif-free rate if the former is larger, and motif-free rate over motif rate otherwise.

### Effect of sequence composition on errors

To investigate whether differences in SMRT sequencing values depend on the presence of different nucleotides, we computed a single nucleotide composition vector *P_Si_* = (*p_A_*, *p_T_*, *p_C_*, *p_G_*) (*p_A_* + *p_T_* + *p_C_* + *p_G_* = 100%), and a dinucleotide composition vector *P_Di_* = (*p_AA_*, *p_AC_*, *p_AG_*, …, *p_TT_*) (*p_AA_* + *p_AC_* + *p_AG_* + … + *p_TT_* = 100%) in each non-repetitive, without fixed differences 100-bp motif-free windows (note: *not* matching motif-free regions; this choice permits to investigate sequence composition effect on the portion of the genome covered by *all* 100-bp motif-free windows). First, we measured the marginal effect of each nucleotide *j=A,C,G,T* on SMRT mismatch error rates as the correlation between *log*(*errorRate*) and *p_j_*. Next, we employed compositional regression models^69,70^ for *log*(*errorRate*) to quantitate the overall effect of single nucleotide and dinucleotide composition on SMRT mismatch error rate. This analysis mirrored the one performed to study the effect of sequence composition on IPD values (see above). The single nucleotide compositional regression model fitted on motif-free windows was then used to predict the SMRT error rates of motif occurrences based on their composition (some motifs are quite short, hence the dinucleotide composition could not be accurately estimated). For each motif type, we computed the differences between these predictions and observed mean IPDs (on logarithmic scale), created their boxplots and performed two-sided t-tests for the mean difference being equal to zero — using a Bonferroni correction to adjust for multiple motif testing (Fig. S17).

### Relationship between errors and IPDs

We considered G4+ and G4-occurrences, as well as non-repetitive, without fixed differences 100-bp motif-free windows (note: *not* matching motif-free regions), in order to obtain three independent groups of regions. 100-bp motif-free windows were randomly subsampled to a number equal to the sum of G4+ and G4-occurrences. We fitted a linear regression model for *log*(*errorRate*) (the SMRT mismatch rates on logarithmic scale) with predictors: (i) residual *log*(*mean IPD*), (ii) region type (G4+ motifs, G4-motifs, baseline motif-free windows), and (iii) interaction between (i) and (ii). The residual *log*(*mean IPD*) in each region was computed as the difference between the observed mean IPD (on logarithmic scale) and the corresponding prediction using single nucleotide composition (compositional regression model fitted on non-repetitive, without fixed differences 100-bp motif-free windows; single nucleotide composition was used because many G4 motifs are short, hence the dinucleotide composition could not be accurately estimated).

### Variants from human-orangutan divergence

We downloaded the 46 species Vertebrate Multiz Alignment^76,77^ from the UCSC Genome Browser (Multiple Alignment Format (MAF) files from https://genome.ucsc.edu/index.html) and considered nucleotide substitutions between human and orangutan. These variants were intersected with our motif occurrences. To obtain an approximate measure of divergence, we divided the number of variants in each motif occurrence by their length. Motifs overlapping with RepeatMasker annotations were excluded also from this analysis.

### Variants from the 1000 Genomes project

We acquired all annotated variants from the 1000 Genomes project (Variant Call Format (VCF) files from http://www.internationalgenome.org/) and intersected the coordinates of those with a global frequency (across all populations) >1% with our motif occurrences. To obtain an approximate measure of diversity, we divided the number of SNPs in each motif by their length. Motifs overlapping with RepeatMasker annotations were excluded also from this analysis.

### Relationship between mutations and IPDs

We compared *log*(*errorRate*) (the SMRT mismatch rates on logarithmic scale) and residual *log*(*mean IPD*) (correcting for single nucleotide composition, see above) between G4+ occurrences with divergence (diversity) levels smaller or equal to the 5^th^ percentile and larger or equal to the 95^th^. Since variants are rare events, a large proportion of motifs have null divergence (diversity), i.e. no genetic variants. As a consequence, there were many more than 5% of the G4+ occurrences with divergence (diversity) equal to 0 (the 5^th^ percentile). In fact, such occurrences were much more abundant than those with divergence (diversity) above the 95^th^ percentile. We subsampled the former 1,000 times to a size equal to the number of the latter. A two-sample, two-sided t-test was performed each time, to test for differences in mean between low and high divergence (diversity) G4+ occurrences. Median p-values (across 1,000 tests) are reported in Table 1, together with median *log*(*errorRate*) and median residual *log*(*meanIP D*) for the same sets of G4+ occurrences.

### Data and Code availability

All scripts are available in public repository https://bitbucket.org/makova-lab/kinetics_wmm. Readers are encouraged to download the latest versions of the scripts directly from the BitBucket repository. The data are available at Data File 1.

## ACKNOWLEDGMENTS

We are grateful to J. Korlach and S. Kingan (Pacific Biosciences Inc.) for comments on the manuscript, to B. Chen, R. Vegesna, F. Cumbo, M. Tomaszkiewicz, and M. Ferguson-Smith for assistance. Funding was provided by the Huck Institutes for the Life Sciences at Penn State and by the Czech Science Foundation (grant 15–02891S).

## AUTHOR CONTRIBUTIONS

WMG, MAC, MC, and RSH performed the computational and statistical analyses, IK and EK performed the biophysical experiments, WMG, MAC, KE, FC, and KDM wrote the manuscript.

## MATERIALS AND CORRESPONDENCE

Correspondence and materials requests should be addressed to KDM or FC.

## REFERENCES

1. Watson, J. D. & Crick, F. H. C. Genetical Implications of the Structure of Deoxyribonucleic Acid. Nature 171, 964–967 (1953).

2. Mirkin, S. M. Discovery of alternative DNA structures: a heroic decade (1979–1989). Front. Biosci. 13, 1064–1071 (2008).

3. Bacolla, A. & Wells, R. D. Non-B DNA conformations, genomic rearrangements, and human disease. J. Biol. Chem. 279, 47411–47414 (2004).

4. Zhao, J., Bacolla, A., Wang, G. & Vasquez, K. M. Non-B DNA structure-induced genetic instability and evolution. Cell. Mol. Life Sci. 67, 43–62 (2010).

5. Maizels, N. G4-associated human diseases. EMBO Rep. e201540607 (2015).

6. Wang, G. & Vasquez, K. M. Z-DNA, an active element in the genome. Front. Biosci. 12, 4424–4438 (2007).

7. Mirkin, S. M. Expandable DNA repeats and human disease. Nature 447, 932–940 (2007).

8. Sen, D. & Gilbert, W. Formation of parallel four-stranded complexes by guanine-rich motifs in DNA and its implications for meiosis. Nature 334, 364–366 (1988).

9. Parkinson, G. N., Lee, M. P. H. & Neidle, S. Crystal structure of parallel quadruplexes from human telomeric DNA. Nature 417, 876–880 (2002).

10. Huppert, J. L. & Balasubramanian, S. Prevalence of quadruplexes in the human genome. Nucleic Acids Res. 33, 2908–2916 (2005).

11. Besnard, E. et al. Unraveling cell type–specific and reprogrammable human replication origin signatures associated with G-quadruplex consensus motifs. Nat. Struct. Mol. Biol. 19, nsmb.2339 (2012).

12. Siddiqui-Jain, A., Grand, C. L., Bearss, D. J. & Hurley, L. H. Direct evidence for a G-quadruplex in a promoter region and its targeting with a small molecule to repress c-MYC transcription. Proc. Natl. Acad. Sci. U. S. A. 99, 11593–11598 (2002).

13. Balasubramanian, S., Hurley, L. H. & Neidle, S. Targeting G-quadruplexes in gene promoters: a novel anticancer strategy? Nat. Rev. Drug Discov. 10, 261–275 (2011).

14. Wittig, B., Dorbic, T. & Rich, A. Transcription is associated with Z-DNA formation in metabolically active permeabilized mammalian cell nuclei. Proc. Natl. Acad. Sci. U. S. A. 88, 2259–2263 (1991).

15. Jansen, A., van der Zande, E., Meert, W., Fink, G. R. & Verstrepen, K. J. Distal chromatin structure influences local nucleosome positions and gene expression. Nucleic Acids Res. 40, 3870–3885 (2012).

16. Belotserkovskii, B. P. et al. Mechanisms and implications of transcription blockage by guanine-rich DNA sequences. Proc. Natl. Acad. Sci. U. S. A. 107, 12816–12821 (2010).

17. Mirkin, S. M. et al. DNA H form requires a homopurine–homopyrimidine mirror repeat. Nature 330, 495–497 (1987).

18. Sawaya, S. et al. Microsatellite tandem repeats are abundant in human promoters and are associated with regulatory elements. PLoS One 8, e54710 (2013).

19. Sinden, R. R., Richard & Sinden, R. Slipped strand DNA structures. Front. Biosci. 12, 4788 (2007).

20. Mirkin, E. V. & Mirkin, S. M. Replication fork stalling at natural impediments. Microbiol. Mol. Biol. Rev. 71, 13–35 (2007).

21. Castel, A. L., Cleary, J. D. & Pearson, C. E. Repeat instability as the basis for human diseases and as a potential target for therapy. Nat. Rev. Mol. Cell Biol. 11, 165–170 (2010).

22. Renton, A. E. et al. A hexanucleotide repeat expansion in C9ORF72 is the cause of chromosome 9p21-linked ALS-FTD. Neuron 72, 257–268 (2011).

23. Haberman, Y., Amariglio, N., Rechavi, G. & Eisenberg, E. Trinucleotide repeats are prevalent among cancer-related genes. Trends Genet. 24, 14–18 (2008).

24. Valton, A.-L. & Prioleau, M.-N. G-Quadruplexes in DNA Replication: A Problem or a Necessity? Trends Genet. 32, 697–706 (2016).

25. Samadashwily, G. M., Raca, G. & Mirkin, S. M. Trinucleotide repeats affect DNA replication in vivo. Nat. Genet. 17, 298–304 (1997).

26. Krasilnikova, M. M. & Mirkin, S. M. Replication stalling at Friedreich’s ataxia (GAA)n repeats in vivo. Mol. Cell. Biol. 24, 2286–2295 (2004).

27. Eddy, S. et al. Evidence for the kinetic partitioning of polymerase activity on G-quadruplex DNA. Biochemistry 54, 3218–3230 (2015).

28. Kang, S., Ohshima, K., Shimizu, M., Amirhaeri, S. & Wells, R. D. Pausing of DNA synthesis in vitro at specific loci in CTG and CGG triplet repeats from human hereditary disease genes. J. Biol. Chem. 270, 27014–27021 (1995).

29. Voineagu, I., Narayanan, V., Lobachev, K. S. & Mirkin, S. M. Replication stalling at unstable inverted repeats: interplay between DNA hairpins and fork stabilizing proteins. Proc. Natl. Acad. Sci. U. S. A. 105, 9936–9941 (2008).

30. Bacolla, A. et al. Breakpoints of gross deletions coincide with non-B DNA conformations. Proc. Natl. Acad. Sci. U. S. A. 101, 14162–14167 (2004).

31. Wang, G., Carbajal, S., Vijg, J., DiGiovanni, J. & Vasquez, K. M. DNA structure-induced genomic instability in vivo. J. Natl. Cancer Inst. 100, 1815–1817 (2008).

32. Bacolla, A. et al. Non-B DNA-forming sequences and WRN deficiency independently increase the frequency of base substitution in human cells. J. Biol. Chem. 286, 10017–10026 (2011).

33. Inagaki, H. et al. Two sequential cleavage reactions on cruciform DNA structures cause palindrome-mediated chromosomal translocations. Nat. Commun. 4, 1592 (2013).

34. Kamat, M. A., Bacolla, A., Cooper, D. N. & Chuzhanova, N. A Role for Non-B DNA Forming Sequences in Mediating Microlesions Causing Human Inherited Disease. Hum. Mutat. 37, 65–73 (2016).

35. Du, X. et al. Potential non-B DNA regions in the human genome are associated with higher rates of nucleotide mutation and expression variation. Nucleic Acids Res. 42, 12367–12379 (2014).

36. Ananda, G. et al. Microsatellite interruptions stabilize primate genomes and exist as population-specific single nucleotide polymorphisms within individual human genomes. PLoS Genet. 10, e1004498 (2014).

37. Eid, J. et al. Real-time DNA sequencing from single polymerase molecules. Science 323, 133–138 (2009).

38. Flusberg, B. A. et al. Direct detection of DNA methylation during single-molecule, real-time sequencing. Nat. Methods 7, 461–465 (2010).

39. Chambers, V. S. et al. High-throughput sequencing of DNA G-quadruplex structures in the human genome. Nat. Biotechnol. 33, 877–881 (2015).

40. Cer, R. Z. et al. Non-B DB v2.0: a database of predicted non-B DNA-forming motifs and its associated tools. Nucleic Acids Res. 41, D94–D100 (2012).

41. Fungtammasan, A. et al. Accurate typing of short tandem repeats from genome-wide sequencing data and its applications. Genome Res. 25, 736–749 (2015).

42. Zook, J. M. et al. Extensive sequencing of seven human genomes to characterize benchmark reference materials. Sci Data 3, 160025 (2016).

43. Cremona, M. A. et al. IWTomics: testing high-resolution sequence-based ‘Omics’ data at multiple locations and scales. Bioinformatics (2018). doi:10.1093/bioinformatics/bty090

44. Nadel, Y., Weisman-Shomer, P. & Fry, M. The fragile X syndrome single strand d(CGG)n nucleotide repeats readily fold back to form unimolecular hairpin structures. J. Biol. Chem. 270, 28970–28977 (1995).

45. Kypr, J., Kejnovská, I., Renciuk, D. & Vorlícková, M. Circular dichroism and conformational polymorphism of DNA. Nucleic Acids Res. 37, 1713–1725 (2009).

46. Neidle, S. & Balasubramanian, S. Quadruplex Nucleic Acids, Royal Society of Chemistry. (2006).

47. Turner, S. et al. Nanoscale apertures having islands of functionality. US Patent (2017).

48. Hodgkinson, A. & Eyre-Walker, A. Variation in the mutation rate across mammalian genomes. Nat. Rev. Genet. 12, 756–766 (2011).

49. Usdin, K. & Woodford, K. J. CGG repeats associated with DNA instability and chromosome fragility form structures that block DNA synthesis in vitro. Nucleic Acids Res. 23, 4202–4209 (1995).

50. Delagoutte, E., Goellner, G. M., Guo, J., Baldacci, G. & McMurray, C. T. Single-stranded DNA-binding protein in vitro eliminates the orientation-dependent impediment to polymerase passage on CAG/CTG repeats. J. Biol. Chem. 283, 13341–13356 (2008).

51. Makova, K. D. & Li, W.-H. Strong male-driven evolution of DNA sequences in humans and apes. Nature 416, 624–626 (2002).

52. 1000 Genomes Project Consortium et al. A global reference for human genetic variation. Nature 526, 68–74 (2015).

53. Locke, D. P. et al. Comparative and demographic analysis of orang-utan genomes. Nature 469, 529–533 (2011).

54. Lander, E. S. et al. Initial sequencing and analysis of the human genome. Nature 409, 860–921 (2001).

55. Schadt, E. E. et al. Modeling kinetic rate variation in third generation DNA sequencing data to detect putative modifications to DNA bases. Genome Res. 23, 129–141 (2013).

56. Sawaya, S., Boocock, J., Black, M. A. & Gemmell, N. J. Exploring possible DNA structures in real-time polymerase kinetics using Pacific Biosciences sequencer data. BMC Bioinformatics 16, 21 (2015).

57. Loomis, E. W. et al. Sequencing the unsequenceable: expanded CGG-repeat alleles of the fragile X gene. Genome Res. 23, 121–128 (2013).

58. Fry, M. & Loeb, L. A. A DNA polymerase alpha pause site is a hot spot for nucleotide misinsertion. Proc. Natl. Acad. Sci. U. S. A. 89, 763–767 (1992).

59. Hile, S. E. & Eckert, K. A. Positive correlation between DNA polymerase alpha-primase pausing and mutagenesis within polypyrimidine/polypurine microsatellite sequences. J. Mol. Biol. 335, 745–759 (2004).

60. Sweasy, J. B., Lauper, J. M. & Eckert, K. A. DNA polymerases and human diseases. Radiat. Res. 166, 693–714 (2006).

61. Makova, K. D. & Hardison, R. C. The effects of chromatin organization on variation in mutation rates in the genome. Nat. Rev. Genet. 16, 213–223 (2015).

62. Cooper, D. N. et al. On the sequence-directed nature of human gene mutation: the role of genomic architecture and the local DNA sequence environment in mediating gene mutations underlying human inherited disease. Hum. Mutat. 32, 1075–1099 (2011).

63. Hänsel-Hertsch, R. et al. G-quadruplex structures mark human regulatory chromatin. Nat. Genet. 48, 1267–1272 (2016).

64. Kouzine, F. et al. Permanganate/S1 Nuclease Footprinting Reveals Non-B DNA Structures with Regulatory Potential across a Mammalian Genome. Cell Syst 4, 344–356.e7 (2017).

65. Simone, R., Fratta, P., Neidle, S., Parkinson, G. N. & Isaacs, A. M. G-quadruplexes: Emerging roles in neurodegenerative diseases and the non-coding transcriptome. FEBS Lett. 589, 1653–1668 (2015).

66. Rhoads, A. & Au, K. F. PacBio Sequencing and Its Applications. Genomics Proteomics Bioinformatics 13, 278–289 (2015).

67. Pini, A. & Vantini, S. Interval-wise testing for functional data. J. Nonparametr. Stat. (2017).

68. Campos-Sánchez, R., Cremona, M. A., Pini, A., Chiaromonte, F. & Makova, K. D. Integration and Fixation Preferences of Human and Mouse Endogenous Retroviruses Uncovered with Functional Data Analysis. PLoS Comput. Biol. 12, e1004956 (2016).

69. Pawlowsky-Glahn, V., Egozcue, J. J. & Tolosana-Delgado, R. Modeling and Analysis of Compositional Data. (John Wiley & Sons, 2015).

70. Aitchison, J. The statistical analysis of compositional data. (Chapman and Hall, 1986).

71. Kejnovská, I. et al. Clustered abasic lesions profoundly change the structure and stability of human telomeric G-quadruplexes. Nucleic Acids Res. 45, 4294–4305 (2017).

72. Dolinnaya, N. G., Ogloblina, A. M. & Yakubovskaya, M. G. Structure, Properties, and Biological Relevance of the DNA and RNA G-Quadruplexes: Overview 50 Years after Their Discovery. Biochemistry 81, 1602–1649 (2016).

73. Dailey, M. M., Miller, M. C., Bates, P. J., Lane, A. N. & Trent, J. O. Resolution and characterization of the structural polymorphism of a single quadruplex-forming sequence. Nucleic Acids Res. 38, 4877–4888 (2010).

74. Taylor, S. & Pollard, K. Hypothesis tests for point-mass mixture data with application to ’omics data with many zero values. Stat. Appl. Genet. Mol. Biol. 8, Article 8 (2009).

75. Lachenbruch, P. A. Analysis of data with clumping at zero. Biom. Z. 18, 351–356 (1976).

76. Blanchette, M. et al. Aligning multiple genomic sequences with the threaded blockset aligner. Genome Res. 14, 708–715 (2004).

77. Harris, R. S. Improved pairwise alignment of genomic DNA. (Pennsylvania State University, 2007).

